# Microfluidic-based mini-metagenomics enables discovery of novel microbial lineages from complex environmental samples

**DOI:** 10.1101/114496

**Authors:** Feiqiao Brian Yu, Paul C. Blainey, Frederik Schulz, Tanja Woyke, Mark A. Horowitz, Stephen R. Quake

## Abstract

Metagenomics and single-cell genomics have enabled the discovery of many new genomes from previously unknown branches of life. However, extracting novel genomes from complex mixtures of metagenomic data can still be challenging and in many respects represents an ill-posed problem which is generally approached with ad hoc methods. Here we present a microfluidic-based mini-metagenomic method which offers a statistically rigorous approach to extract novel microbial genomes from complex samples. In addition, by generating 96 sub-samples from each environmental sample, this method maintains high throughput, reduces sample complexity, and preserves single-cell resolution. We used this approach to analyze two hot spring samples from Yellowstone National Park and extracted 29 new genomes larger than 0.5 Mbps. These genomes represent novel lineages at different taxonomic levels, including three deeply branching lineages. Functional analysis revealed that these organisms utilize diverse pathways for energy metabolism. The resolution of this mini-metagenomic method enabled accurate quantification of genome abundance, even for genomes less than 1% in relative abundance. Our analyses also revealed a wide range of genome level single nucleotide polymorphism (SNP) distributions with nonsynonymous to synonymous ratio indicative of low to moderate environmental selection. The scale, resolution, and statistical power of microfluidic-based mini-metagenomic make it a powerful tool to dissect the genomic structure microbial communities while effectively preserving the fundamental unit of biology, the single cell.

## Main Text

Advances in sequencing technologies have enabled the development of shotgun and single-cell metagenomic approaches to investigate environmental microbial communities. These studies revealed many previously uncharacterized genomes^1-4^, increasing the total number of sequenced microbial genomes to more than 50,000 (Joint Genome Institute’s Integrated Microbial Genomes database, accessed December 1, 2016). However, the majority of environmental microbial diversity remains uncharacterized due to limitations in current techniques. Conventional shotgun metagenomic sequencing offers the ability to assemble genomes from heterogeneous samples, but are effective only if the complexity of the sample is not too great^5^. Furthermore, it is often difficult to separate contigs belonging to closely related organisms because techniques designed to resolve these differences, such as tetranucleotide analysis, depend on ad hoc assumptions about nucleotide usage^6^. It is possible to perform rigorous assemblies from independent single-cell genome amplifications but at the expense of lowering throughput^7,8^. In addition, when performed in plates, single-cell sequencing approaches are typically expensive and laborious, although microfluidic approaches have alleviated that limitation^9^.

We developed a new mini-metagenomics approach that combines the advantages of shotgun and single-cell metagenomic analyses. We used microfluidic parallelization to separate an environmental sample into many small sub-samples containing 5-10 cells, significantly reducing complexity of each sub-sample and allowing high quality assembly while enabling higher throughput than typical single-cell methods. Although each sub-sample contains a limited mixture of several genomes, we regain single-cell resolution through correlations of genome co-occurrence across sub-samples, which results in a rigorous statistical interpretation of confidence and genome association.

We validated this approach using a synthetic mixture of defined microbial species and then applied it to analyze two hot spring samples from Yellowstone National Park. Among 29 genomes larger than 0.5 Mbps, most belong to known bacterial and archaeal phyla but represent novel lineages at lower taxonomic levels. We identified three genomes representing deeply branching novel phylogenies. Functional analysis revealed different metabolic pathways that cells may use to achieve the same biochemical process such as nitrogen and sulfur reduction. Using information associated with genome occurrence across sub-samples, we further assessed abundance and genome variations at the single-cell level. Our analyses demonstrate the power of the mini-metagenomic approach in de-convolving genomes from complex samples and assessing diversity in a mixed microbial population.

Microfluidic-based mini-metagenomics begins with microfluidic partitioning of each environmental sample randomly into 96 sub-samples with 5-10 cells per sub-sample (Fig. 1A). Lysis and MDA (Multiple Displacement Amplification) are performed in independent chambers of an automated and commercially available Fluidigm C1 Integrated Fluidic Circuit (IFC) (Fig. 1B and Fig. S1A), which significantly reduces the time and effort required to perform these reactions. After whole-genome amplification, sequencing libraries are prepared with distinct barcodes labeling DNA derived from each sub-sample and sequenced on the Illumina Nextseq platform (Fig. 1C). Because of the small volumes used for microfluidic reactions, precious microbial samples and/or low cell concentrations that may not yield enough DNA for shotgun metagenomics can still be analyzed using our process. Another advantage of using a microfluidic platform is the enclosed reaction environments that limit potential contamination often observed with low input MDA reactions in well plates, a problem that, with other approaches, requires sophisticated protocols for contamination prevention and removal^10^. In addition, smaller MDA reaction volumes (~300 nL) are associated with lower gain and less amplification bias, which effectively improves coverage uniformity of amplified genomes^11^ (Methods).

**Fig. 1.**
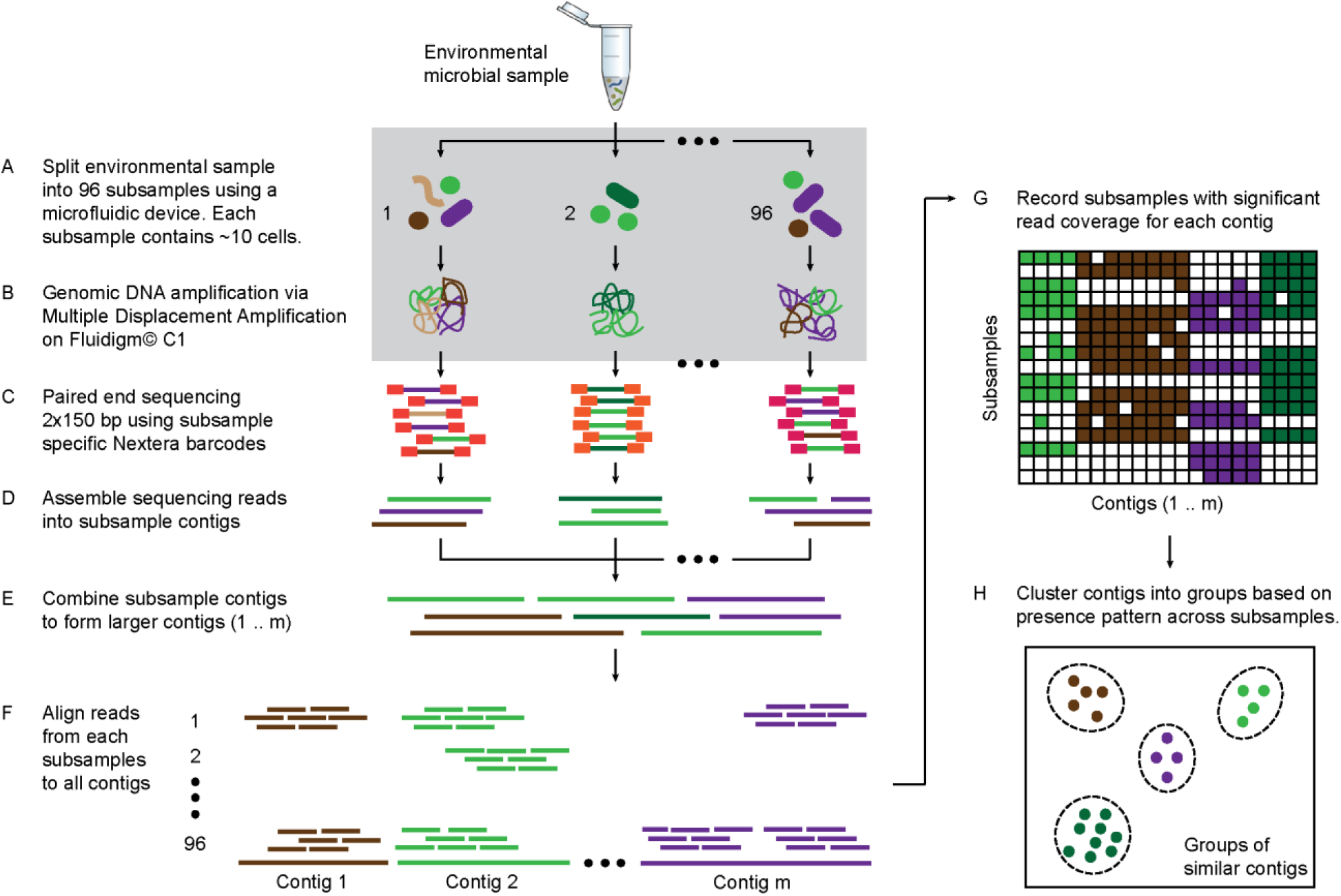
Microfluidic-based mini-metagenomics pipeline. (A) An environmental microbial sample is loaded onto a Fluidigm C1 IFC at the appropriate concentration so that cells are randomly dispersed into 96 microfluidic chambers at 5-10 cells per chamber. (B) Lysis and MDA are performed on the microfluidic device to generate 1–100 ng genomic DNA per sub-sample. (C) Nextera libraries are prepared from the amplified DNA off-chip and sequenced using 2×150 bp runs on the Illumina NextSeq platform. (D) Sequencing reads from each sub-sample are first assembled independently, then (E) sub-sample contigs are combined to form longer mini-metagenomic contigs. Only contigs longer than 10 kbp are processed in the following steps. (F) Reads from each sub-sample are aligned to mini-metagenomic contigs >10 kbp. (G) An occurrence map is generated, demonstrating the presence pattern of each contig in all sub-samples based on coverage. (H) Finally, contigs are binned into genomes clusters based on a pairwise *p* value generated from co-occurrence information.

Sequence data was processed through a custom bioinformatics pipeline that takes advantage of information encapsulated in distinct yet related sub-samples. Reads from each sub-sample were trimmed and assembled into sub-sample contigs (Fig. 1D), creating genome subassemblies. Since cells representing the same genome may appear in multiple sub-samples, overlapping genome subassemblies can be identified. Therefore, combined assembly of sub-sample reads and contigs results in longer mini-metagenomic contigs from which more meaningful biological information can be inferred (Fig. 1E). With this mini-metagenomic approach, the presence of cells from the same phylogenetic group which are randomly partitioned into different sub-samples provides a physically defined approach to bin metagenomic contigs. Aligning reads from each sub-sample to mini-metagenomic contigs enabled us to determine the co-occurrence pattern of each contig (Fig. 1F, G). Based on presence patterns across sub-samples, a *p* value for each pair of contigs was computed based on Fisher’s exact test (Methods), where a small *p* was interpreted as an indicator that two contigs belonging to cells of the same genome. Finally, co-occurrence based *p* values was used as a pairwise distance metric for sequence-independent contig (Fig. 1H, Methods).

We tested performance of the mini-metagenomic amplification on the Fluidigm C1 IFC using a mixture of five bacterial species with known genomes (*E. vietnamensis, S. oneidensis, E. coli, M. ruber,* and *P. putida*) (table S1, Methods). We first used a dilution of the control sample at ~10 cells per sub-sample. Then, we further diluted the control sample so that each sub-sample contains 0.5 cells on average, effectively performing microfluidic single-cell MDA with the same downstream steps (Methods). Performing MDA from multiple cells in a 300 nL microfluidic chamber improves genome coverage at similar sequencing depths because smaller MDA reaction volumes tend to reduce amplification gain and bias^11,12^.

Thirty mini-metagenomic and 36 single-cell limiting dilution sub-samples yielded 169 and 194 million paired end reads, respectively. Initial trimming removed 14±1.1% (s.d.) reads; 80±1.5% (s.d.) of the remaining reads mapped uniquely to reference genomes (Fig. S2A). Unmapped reads made up only 3% of all reads (Fig. S2B). Improperly mapped reads (3±0.9% s.d.) were mostly short, low quality sequences. Chimeras represented <1% of the reads. Mini-metagenomic MDA reactions produce higher median coverage than single-cell MDA reactions at all sequencing depths (Fig. S3A). The proportion of assembled genome as a function of aligned genome, however, did not differ significantly between mini-metagenomic and single-cell methods (Fig. S3B). Therefore, data from mock bacterial communities demonstrate that with the mini-metagenomic method, less sequencing cost is required to recover similar genome coverage as compared to single cell experiments, with the improvement mostly associated with the amplification rather than assembly steps.

Next, we performed microfluidic mini-metagenomic sequencing on two hot spring samples from Yellowstone National Park (Methods). Sample #1 was collected from Bijah Spring and sample #2 was collected from Mound Spring (table S3). 121 and 133 million paired end reads were obtained from the two samples. We removed sub-samples with less than 800,000 paired end reads, yielding 49 and 93 sub-samples. After quality filtering, >90% reads from each sub-sample were incorporated into contigs during independent assemblies (Fig. S4). Re-assembling combined sub-sample reads increased contig length (Fig. S5A). We obtained 643 and 1474 contigs longer than 10 kbp from the two samples, respectively, and used these contigs for subsequent analyses (Fig. S5B). To compare this performance to shotgun metagenomic assemblies, we obtained 32.5 million and 51.4 million shotgun metagenomic reads from Bijah Spring and Mound Spring samples and down sampled combined mini-metagenomic reads to match shotgun sequencing depths. After assembly (Methods), we observed that for the Bijah Spring sample, where 49 sub-samples are available, the shotgun metagenomic assembly produced more contigs over 10 kbp (Fig. S6A). However, for the Mound Spring sample, where 93 sub-samples were used, the mini-metagenomic assembly produced more contigs over 10 kbp (Fig. S6A). The largest contig from the mini-metagenomic assembly was also longer for the Mound Spring sample (Fig. S6B). Therefore, it is likely that increasing the number of mini-metagenomic sub-samples improve assembly performance compared to shotgun metagenomics. It is also possible that the observed assembly improvement is due to higher complexity of the Mound Spring bacterial community, validating our hypothesis that the mini-metagenomic method benefits the analysis of complex microbial populations.

Another advantage of mini-metagenomics lies in the information from sub-samples. In order to bin contigs into genomes, reads from each sub-sample were aligned back to mini-metagenomic contigs and coverage was tabulated (Fig. 2A, B and Fig. S5C, D). Many contig sets share similar coverage patterns across sub-samples, suggesting that they originate from the same genome. Because MDA results in variable coverage profiles, we generated a binary occurrence map by applying a coverage threshold (Methods). Pairwise *p* values were computed with Fisher’s exact test (Fig. 2 and Fig. S7). Finally, dimensionality reduction using pairwise *p* values as a distance metric generated clusters of contigs belonging to the same genomes^13^ (Fig. S8).

**Fig. 2.**
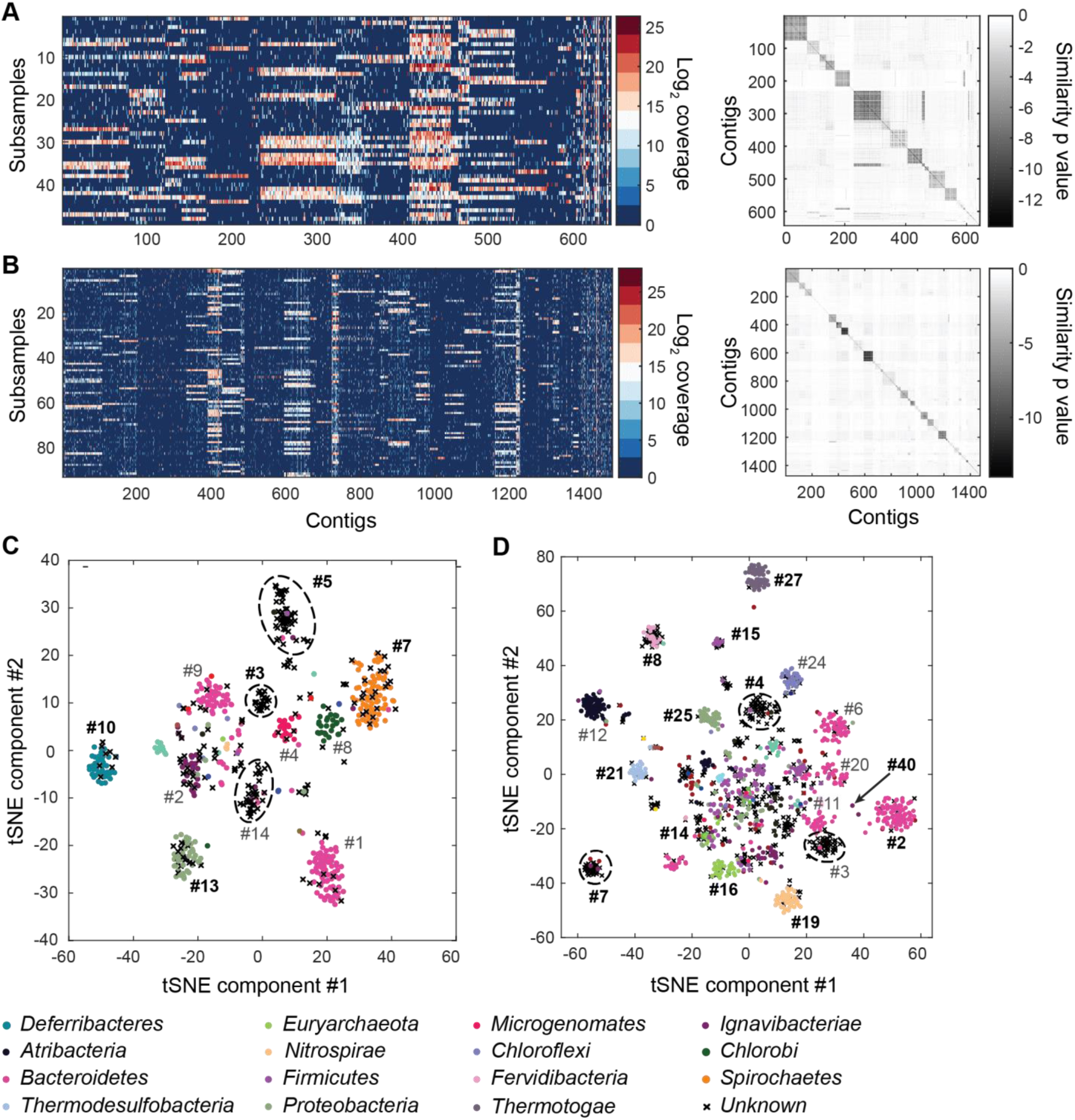
Genomic bins extracted from microfluidic-based mini-metagenomic sequencing of Yellowstone National Park samples. Two samples from Bijah Spring in Mammoth Norris Corridor (A, C) and Mound Spring in Lower Geyser Spring Basin (B, D) are collected from Yellowstone National Park and analyzed using the microfluidic-based mini-metagenomic pipeline. (A, B) Heat maps of contig coverage across sub-samples are clustered hierarchically to reveal contigs that appear in similar sets of sub-samples (left). Colors represent logarithm of coverage in terms of number of base pairs in base 2. Pairwise *p* values generated using Fisher’s exact test based on co-occurrence pattern of contig pairs reveal contig clusters (right). Shading here represents logarithm of *p* value in base 10 after correcting for multiple comparisons. (C, D) tSNE dimensionality reduction generated from pairwise *p* values. Each point represents a 10 kbp or longer contig. Colors represent assignment of each contig to a particular phylum based on annotation of genes on the contig. Black X’s represent contigs unable to be assigned to any phylum because too many genes have unknown annotation. Genome bins larger than 0.5 Mbp are numbered and those with substantial numbers of single-copy marker genes for incorporation into Figure 4 are labeled in bold. Dotted circles outline genomes predominantly containing contigs unassigned at the phylum level.

To verify the validity of presence-based contig clusters, we annotated all predicted open reading frames (ORFs)^14^ (Methods). The relationship between contig length and the number of genes found is linear (Fig. S9), consistent with small non-coding regions in bacterial genomes. Annotations were found for ~50% of the genes. Based on gene annotations from the same contig, lineage assignment was performed. If protein sequences of most genes on a contig were distantly related to known sequences, the contig was designated as unknown (Fig. 2C, D). Approximately 70% of all contigs were assigned at the phylum level. Contigs with known assignment and from the same cluster always belonged to the same phylum, demonstrating that the presence-based method can correctly bin metagenomic contigs into genomes (Fig. 2C, D). Some unassigned contigs are scattered among genome bins with known phylum level assignments. Unassigned contigs likely represent novel genes. There are also genome bins containing predominantly unassigned contigs indicated by dotted circles (Fig. 2C, D), likely representing deeply branching lineages.

We selected 29 partial genome bins from both hot spring samples with genome sizes over 0.5 Mbp for downstream analyses (Fig. 3A). Assessment of genome bins demonstrates various levels of completeness and <5% marker gene duplication (Fig. 3B, Fig. S10). Genome completeness is not necessarily correlated with genome size because completeness is assessed through single-copy marker genes^15^. Using lineage assignment based on gene annotations, we identified eight genome bins with known phylum level assignments from the Bijah Spring sample. Three genomes (Bijah #3, Bijah #5, and Bijah #14) had unknown assignments (Fig. 2C). In the Mound Spring sample, more reads and sub-samples resulted in 14 genomes with known phylum level assignments and three unassigned genomes (Mound #3, Mound #4, and Mound #7) were extracted (Fig. 2D). A singleton contig 3.3 Mbp in length (#40) was also included as a separate genome. In both metagenomic samples, the ability to separate distinct clusters of *Bacteroidetes* contigs represents an advantage of our sequence-independent binning approach, where the presence of bacterial species across microfluidic chambers is determined only by Poisson distribution. Hence, the likelihood that closely related bacterial cells are always isolated into the same chambers is small.

**Fig. 3.**
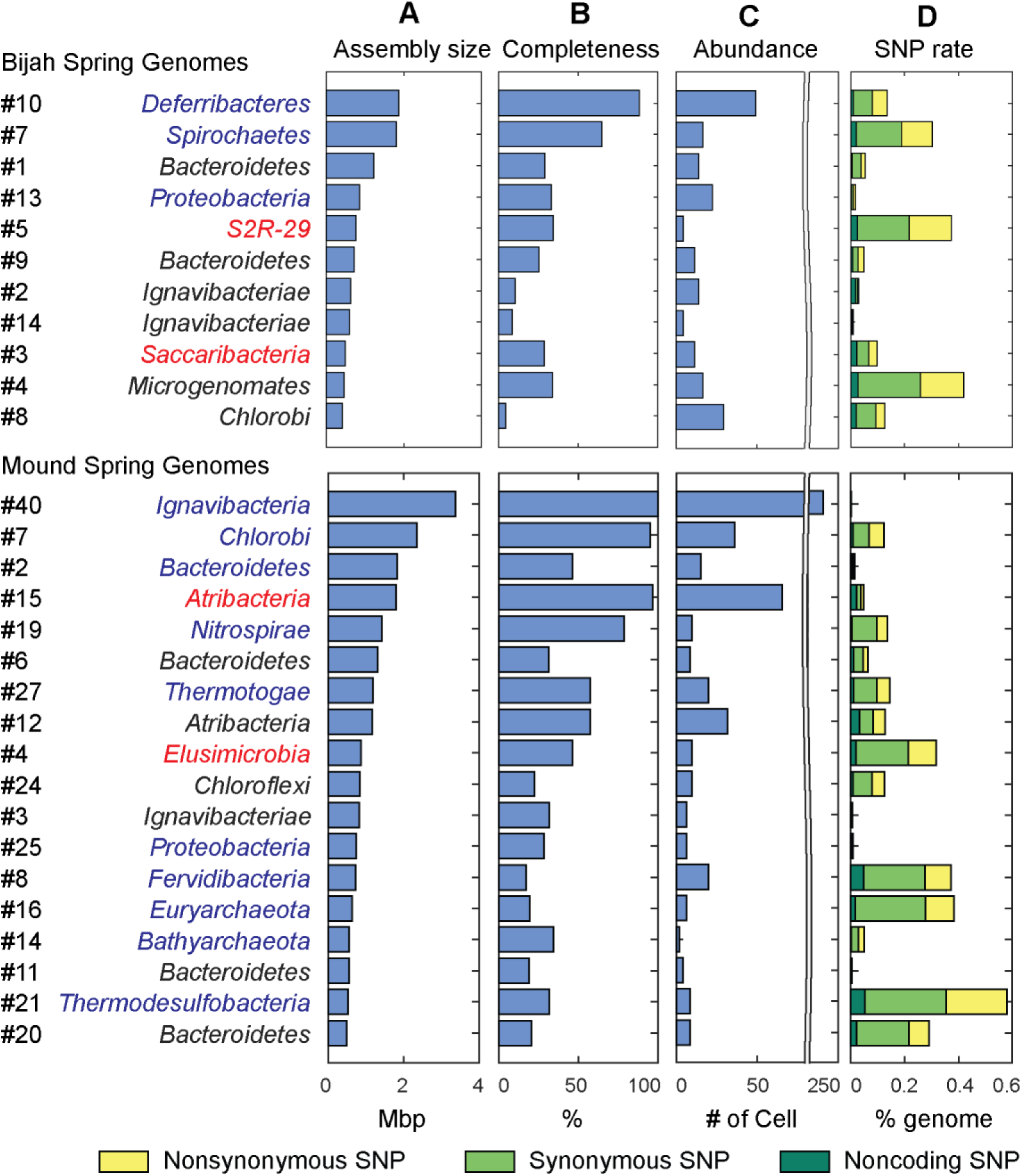
Genome statistics, abundance, and variation. Genomes of Bijah Spring and Mound Spring samples are sorted by genome size (A). Names represent phylum level assignment based on annotated genes (Fig. 2), concatenated marker gene phylogenetic tree (Fig. 4), or individual marker gene trees (Fig. S10, S11). (B) Genome completeness is assessed through single-copy marker genes; those incorporated into Figure 4 have phyla names colored in blue (for short branching lineages) or red (for deep branching lineages). (C) Abundance is derived from the occurrence pattern of contig clusters, where Poisson distribution is used to infer number of cells processed. (D) SNPs are tabulated and normalized by total size of sequenced genome. Most SNPs are in coding regions of the genome, of which the majority are synonymous.

Since extracted genomes of known phyla often represent novel lineages at lower taxonomic levels, we attempt to identify their phylogenetic placements. Seventeen genomes (shown in bold in Fig. 2C, D), including four unassigned genomes, were complete enough for phylogenetic tree construction based on 56 single copy marker genes^2^ (Fig. 4; Methods). All marker gene based phylogenetic assignments were consistent with annotation based assignments. We identified 13 genomes with short to medium branch length from known lineages (red), and three deeply branching lineages that may represent potentially novel phyla (red and starred). In additional to bacterial lineages, we also extracted partial archaeal genomes belonging to *Euryarchaeota* (Mound #16) and *Bathyarchaeota* (Mound #14) (Fig. 4). The remaining two genomes (Bijah #14 and Mound #3) with unassigned phylogeny that were also not complete enough for incorporation into the final phylogenetic tree were examined independently. Phylogenetic trees built from blast results of individual ribosomal protein sequences suggest that both genomes belong to the phylum *Ignavibacteriae* (Fig. S11, S12).

**Fig. 4.**
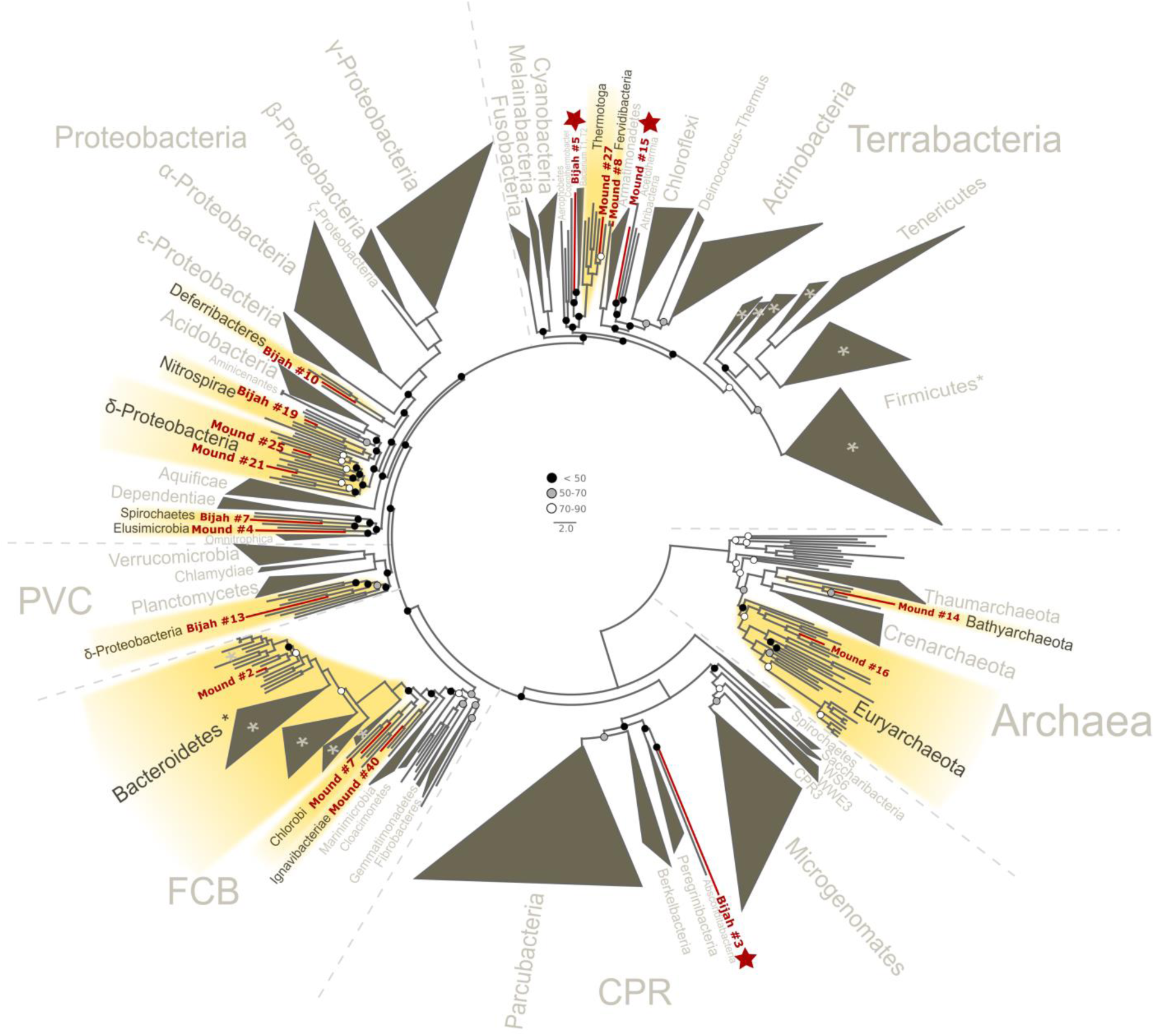
Phylogenetic distribution of selected Yellowstone hot spring genomes (red branches) across a representative set of bacterial and archaeal lineages. Query genomes which potentially represent novel phyla are marked with a star, and those falling into known phyla are highlighted in yellow. Query genomes with less than 28 out of 56 marker proteins and/or less than 30% of informative positions in the sequence alignment are indicated by a filled red circle. Bootstrap support values are displayed at the nodes as filled circles in the following categories: no support (black), weak support (grey), moderate support (white), while absence of circles indicates strong support (>90 bootstrap support). For details on taxon sampling and tree inference, see Methods.

To understand more about the metabolism of these organisms, we performed BLAST of all ORFs contained in each genome against the KEGG database and mapped results onto KEGG modules (Methods). Figure S12 illustrates the proportion of identified KEGG module genes from each genome belonging to pathways associated with nitrogen, methane, and sulfur metabolism. At the community level, the Mound Spring population displays more potential for methanogenesis than the Bijah Spring population, and methanogenesis can be carried out by members of the *Euryarchaeota* (Mound # 16) lineages involving *mcr* and *hdr* complex^16^. Formaldehyde assimilation, on the other hand, can be carried out in both communities. In nitrogen metabolism, we identified five genomes (Mound #40 #2 #19 #21 and Bijah #2) carrying *nar, nrf,* and *nir* genes, possibly participating in the conversion of nitrate to ammonia^17-19^. Interestingly, the archaeal genome belonging to *Euryarchaeota* (Mound #16) carries *nifD* and *nifH,* capable of converting nitrogen directly to ammonia^20^ (Fig. 5B). Although we did not identify denitrification genes in the Mound Spring community, a genome extracted from Bijah Spring belonging to *Spirochaetes* carries *nirS, norBC,* and *nosZ* genes, capable of reducing nitrite to nitrogen (Fig. 5A).

**Fig. 5.**
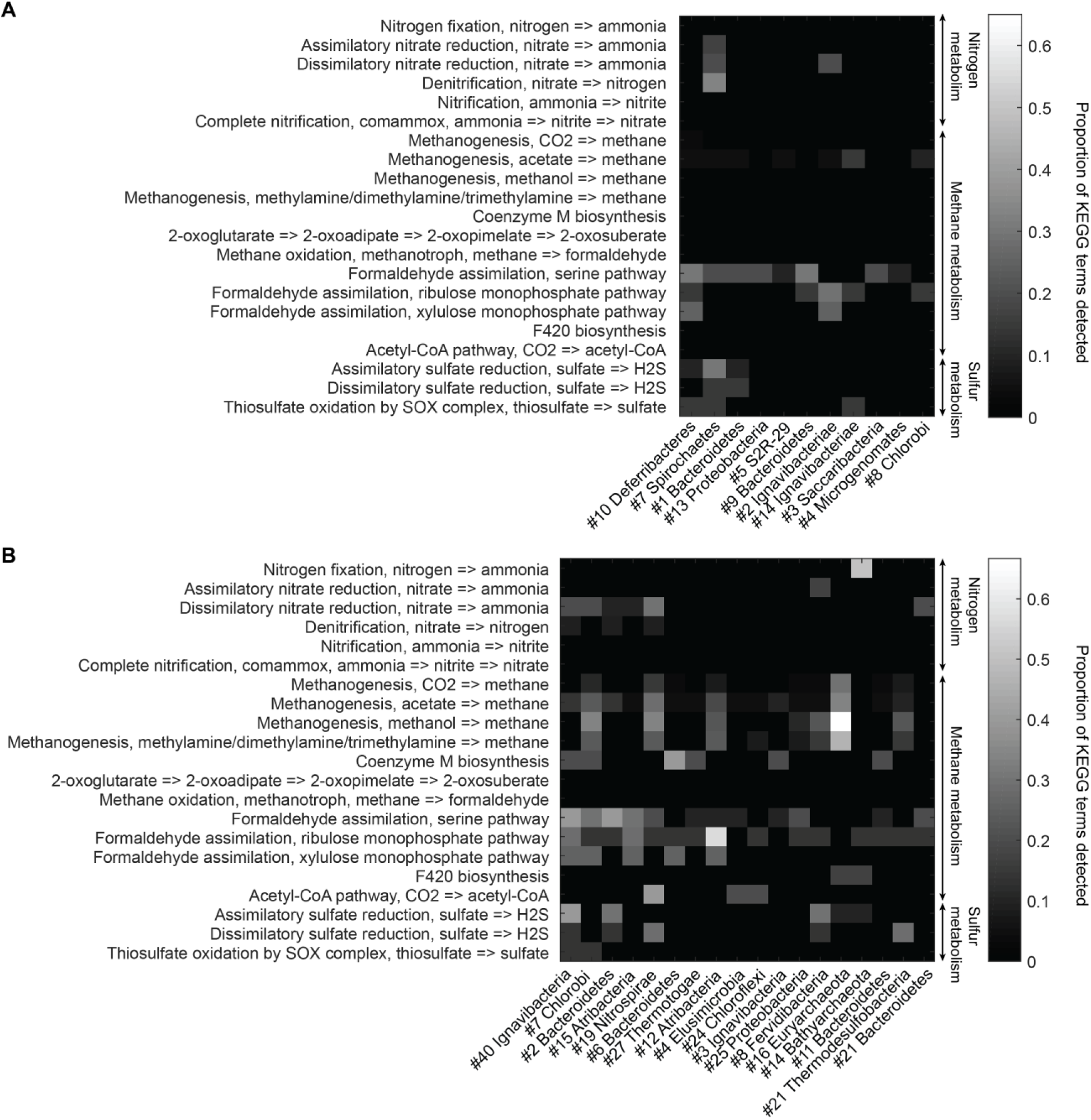
Abundant genes involved in energy metabolism of (A) Bijah Spring and (B) Mound Spring genomes. Each row represents description of a pathway based on KEGG energy metabolism modules. Each column represents one genome bin. Shading of each square represents the ratio of genes in each KEGG module that are also present in a particular genome bin. Modules are labeled as nitrogen metabolism, methane metabolism, or sulfur metabolism.

Bacterial clades associated with thermophilic environments in the Yellowstone National Park are known for their sulfur reduction (converting sulfate to sulfide) activities^21,22^. Two well characterized pathways are assimilatory and dissimilatory processes, where sulfur is either incorporated into cellular materials or serve only as the terminal electron acceptor^23^. We identified genes associated with the assimilatory sulfate reduction pathway (*cysNC, sat, cysC, cysH,* and *cysI*) from three Mound Spring genomes (Mound #40 #2 #8) and one Bijah Spring genome (Bijah #7). In the Mound Spring population, we also identified key genes (*dsrAB*) responsible for dissimilartory sulfate reduction from *Nitrospirae* (Mound #19) and *Termodesulfobacteria* (Mound #21) genomes (Fig. 5). These insights based on genes with known annotations demonstrate the ability of mini-metagenomics to reveal different metabolic pathways associated with individual genomes within and across environmental samples.

The presence of related genome sequence across sub-samples can be used to estimate genome abundance through counting cells, which is not affected by amplification bias.

Assuming a Poisson distribution, we quantified the number of detected cells of all genomes in both samples (Fig. 3C, Fig. S13; Methods). Assembled genomes from the Bijah sample covered one order of magnitude in abundance. Genomes #10 (*Deferribacteres*) and #8 (*Chlorobi*) are most abundant, with 49 and 29 cells represented. On the lower end, Bijah #5 (*S2R-29*) is only represented in four sub-samples, most likely representing only four cells (Fig. S13A). In the Mound Spring sample, more sub-samples were sequenced, allowing the quantification for a higher range of abundance across two orders of magnitude (Fig. S13B). Mound #40 (*Ignavibacteria*) is the most abundant lineage, appearing in all but seven sub-samples. The abundance of this genome is likely the reason for the successful assembly of its entire genome (Fig. 3D). Mound #7 (*Chlorobi*) and deeply branching Mound #15 were also abundant, allowing the capture of 36 and 65 cells respectively. For rare organisms, we detected Mound #11 (*Bacteroidetes*) and Mound #14 (*Bathyarchaeota*), which were present at < 1%. In total, examining genomes larger than 0.5 Mbp, we captured 192 and 509 cells from Bijah and Mound Springs, respectively, corresponding to four to six cells per sub-sample. Even though this number is smaller than our intended ten cells per chamber, the difference is likely due to lysis inefficiency or smaller genomes excluded in our analysis.

In addition to quantifying abundance through counting single cells, genetic variations among individual cells with the same genome can be assessed as well. We quantify observed single nucleotide polymorphisms (SNPs) in each assembled genome normalized to the total assembled genome size (Fig. 3D, Methods). We found a wide distribution in SNP abundance among phylogenies. All *Ignavibacteriae* and three *Bacteroidetes* genomes had low SNP rates < 1%. In the Bijah Spring sample, members of *Spirochaetes, Microgenomates,* and *S2R-29* contained the most SNPs at 0.3 – 0.4% of the assembled genome. In the Mound Spring sample, members of *Thermodesulfobacteria* had most SNPs (0.6%). Several other groups belonging to *Bacteroidetes, Fervidibacteria, Euryarchaeota,* and deeply branching Mound #4 all have 0.3 – 0.4% SNP rate. It is not the case that larger, more complete, or more abundant genomes contain more SNPs in a population (Fig. S14), indicating that the observed SNP rate is a biological property of the particular genome. The ratio of nonsynonymous to synonymous SNP rates (dN/dS) is typically a measure of the strength of environmental selection^24^, with one indicating no selection and zero indicating strong selection. Among diverse genomes that allow more accurate assessment of SNP ratios, we found dN/dS ranging from 0.4 – 0.9 in the hot spring genomes, illustrating that different phylogenies in the same hot spring environment may be subjected to different weak to moderate selection pressures. Taken together, the microfluidic-base mini-metagenomic method leverages the statistical power from multiple sub-samples to more effectively bin contigs for functional analysis, quantify abundance, and assess genomic variation.

Previous work has implemented similar concepts of sequencing multiple bacterial cells together in order to increase throughput. However, the authors treated each sub-sample as an independent entity throughout their bioinformatic analysis^25^. The novelty of our analysis derives from treating information from sub-samples as different but overlapping sections of a more complex metagenome. Such an approach echoes recent works that use coverage from many shotgun metagenomic samples containing a similar set of microbial phylogenies but with different abundances in order to aid bioinformatic genome binning^26,27^. Our approach differs with these methods in several aspects. First, our approach generates sub-samples from one, possibly low volume, microbial sample, eliminating the need to collect multiple spatial or temporal metagenomic samples. In addition, because of low complexity, each sub-sample does not need to be sequenced as deeply as using shotgun metagenomic methods, significantly saving sequencing cost. Because of MDA bias, we do not use coverage depth directly. Instead, thresholding coverage depth into a presence profile can be seen as digitizing an otherwise noisy analog signal. Such digitization reduces sensitivity to amplification bias associated with analog coverage signals.

Microfluidic-based mini-metagenomics provide researchers with two independent experimental design parameters, namely the average number of cells per microfluidic chamber and the number of chambers to sequence, in order to optimize the protocol for microbial samples of different complexities. The statistical power of the presence based binning technique benefits from both the presence and absence of genomes across sub-samples. Therefore, for less complex samples, reducing the number of cells per microfluidic chamber would ensure cells of the same lineage do not appear in all sub-samples. For more complex samples, the number of cells per chamber can be increased to capture higher diversity. One can even carry out multiple runs with different loading densities to tackle both abundant and rare species. More importantly, because microfluidic-based mini-metagenomics provide orthogonal information to tetranucleotide or coverage information derived from traditional metagenomic sequencing, combining these approaches would represent exciting future directions for deriving a more comprehensive picture of the microbial world.

## Methods

### Mock sample construction

The artificial bacterial community used to test the mini-metagenomic approach was constructed using five model species with different GC content provided by the Joint Genome Institute (table S1). Each species was cultured independently, quantified via platting, and combined to create an artificial mixed population. Ultrapure glycerol (Invitrogen) was immediately added to the mixture at 30% and stored at −80°C.

### Environmental sample collection and storage

The environmental samples used in this study were collected from two separate hot springs from the Yellowstone National Park (table S2). Sample #1 was collected from sediments of the Bijah Spring in the Mammoth Norris Corridor area. Sample #2 was collected from sediment near Mound Spring in the Lower Geyser Basin region. Samples were spaced in 2 mL tubes and soaked in 50% ethanol onsite. Upon returning, samples were transferred to −80°C for long term storage.

### Sample preparation and dilution for mini-metagenomic pipeline

Each mock sample was thawed on ice and centrifuged at 5000 x g for 10 min at room temperature. Supernatant was removed and cells were re-suspended in 1% NaCl. Each sample from Yellowstone was also thawed on ice. The tube is vortexed briefly to suspend cells but not large particles and debris. 1 mL of sample from the top of the tube was removed, placed in a new 1.5 mL tube, and spun down at 5000 × g for 10 minutes to pellet the cells. Supernatant was removed and cells were re-suspended in 1% NaCl. After resuspension, microscopy was performed to quantify cell concentration. Each sample was then diluted in 1% NaCl or PBS to a final concentration of ~2×10^6^ cell/mL, corresponding to ~10 cells per chamber on the Fluidigm C1 microfluidic IFC (Integrated Fluidic Circuit).

### Microfluidic genomic DNA amplification on Fluidigm C1 auto prep system

Because of the small amount of DNA associated with 5-10 cells, DNA contamination is a concern for MDA reactions. To reduce DNA contamination, we treated the C1 microfluidic chip, all tubes, and buffers under UV (Strategene) irradiation for 30 minutes following suggestions of Woyke et al^10^. Reagents containing enzymes, DNA oligonucleotides, or dNTPs were not treated. After UV treatment, the C1 IFC was primed following standard protocol (Fluidigm). The diluted bacterial sample was loaded onto the chip using a modified version of the loading protocol where washing was not performed, as the capture sites were too large for bacterial cells. Hence, they act essentially as chambers into which cells were randomly dispersed. Following cell loading, whole genome amplification via MDA was performed on-chip in 96 independent reactions. A lysozyme (Epicenter) digestion step was added before alkaline denaturation of DNA. After alkaline denaturation of DNA, neutralization and MDA were performed (Qiagen REPLI-g single cell kit). Concentrations of all reagents were adjusted to match the 384 well plate-based protocol developed by the single-cell group at DOE’s Joint Genome Institute but adapted for volumes of the Fluidigm C1 IFC ^28^. Lysozyme digest was performed at 37 °C for 30 min, alkaline denaturation for 10 min at 65 °C, and MDA for 2 hours and 45 minutes at 30 °C. The detailed custom C1 scripts and reagent compositions will be made available through Fluidigm’s ScriptHub.

### Library preparation and sequencing

Amplified genomic DNA from all sub-samples was harvested into a 96 well plate. The concentration of each sub-sample is quantified independently using the high sensitivity large fragment analysis kit (AATI). Following quantification, DNA from each sub-sample is diluted to 0.1 − 0.3 ng/μL, the input range of the Nextera XT library prep pipeline. Nextera XT V2 libraries (Illumina) are made with dual sequencing indices, pooled, and purified with 0.75 volumes of AMpure beads (Agencourt). Illumina Nextseq (Illumina) 2×150 bp sequencing runs are performed on each library pool.

### Contig construction

Sequencing reads were filtered with Trimmomatic V0.30 in paired end mode with options “ILLUMINACUP:adapters.fa:3:30:10:3:TRUE SLIDINGWINDOW:10:25 MAXINFO:120:0.3 LEADING:30 TRAILING:30 MINLEN:30” to remove possible occurrences of Nextera indices and low quality bases^29^. Filtered reads from each sub-sample were clustered using DNACLUST, with k=5 and a similarity threshold of 0.98, in order to remove reads from highly covered regions^30^. Then, assembly was performed using SPAdes V3.5.0 with the sc and careful flags asserted^31^. From all sub-sample assembly output, corrected reads were extracted and combined. The combined corrected reads from all sub-samples are assembled again via SPAdes V3.5.0 with kmer values of 33,55,77,99. Finally contigs longer than 10 kbp are retained for downstream analyses.

### Gene annotation

Contigs were uploaded to JGI’s Integrated Microbial Genomes’s Expert Review online database (IMG/ER). Annotated is performed via IMG/ER^14^. Briefly, structural annotations are performed to identify CRISPRs (pilercr), tRNA (tRNAscan), and rRNA (hmmsearch). Protein coding genes are identified with a set of 4 *ab initio* gene prediction tools: GeneMark, Prodigal, MetaGeneAnnotator, and FragGeneScan. Finally, functional annotation is achieved by associating protein coding genes with COGs, Pfams, KO terms, EC numbers. Phylogenetic lineage was assigned to each contig based on gene assignment.

#### Contig co-occurrence distance score and binning

Corrected reads from each sub-sample were aligned back to assembled contigs over 10 kbp using Bowtie2 V2.2.6 with options “--very-sensitive-local-I 0 −X 1000” ^32^. Total coverage in terms of number of base pairs covered for every contig from every sub-sample was tabulated and subjected to a log transform in base two (Fig. 2A, B). A threshold value of 2^11^ is used to determine if a contig had significant presence in a sub-sample. The sensitivity of the contig binning results to this threshold was low, with threshold values of 2^9^–2^13^ producing similar results. After thresholding contig coverage, the occurrence pattern of all contigs across sub-samples was obtained. Based on co-occurrence patterns, a confidence score for each pair of contigs can be computed based on Fisher’s exact test (Fig. 2A, B). This score represented the probability of incorrectly rejecting the null hypothesis that two contigs display a particular co-occurrence pattern by chance and can be interpreted as the likelihood of two contigs belonging to cells of the same genome, with lower values increasing this likelihood. Below, we demonstrate an example of how to calculate a similarity *p* value based on co-occurrence pattern of two contigs across a set of sub-samples in Figure S7. For every pair of contigs X and Y, the null hypothesis states that their co-occurrence patterns are not correlated with each other. Then, we tabulate 4 values A, B, C, D as shown below.

A = number of sub-samples where X is present and Y is present.

B = number of sub-samples where X is absent and Y is present.

C = number of sub-samples where X is present and Y is absent.

D = number of sub-samples where X is absent and Y is absent.

To test the null hypothesis, we compute a *p* value according to the following equation

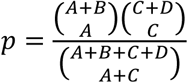

From the simplified example, we see that X1 and Y1 are both present in 8 sub-samples, hence A=8. In addition, B=1, C=2, D=6, yielding *p* = 0.013 (Fig. S7A). Since *p* is small, we reject the null hypothesis and conclude that X1 and Y1 are correlated. On the other hand, co-occurrence patterns of X2 and Y2 produces p=0.363, which is not small enough to reject the null hypothesis (Fig. S7B).

Tabulating all pairwise *p* values, we created a similarity matrix containing values between 0 and 1, which can also act as a pairwise distance metric. Using this pairwise distance metric, we performed dimensionality reduction of all contigs using tSNE^13^ (Fig. 2C, D, Fig. S7).

### Phylogenetic lineage construction

A set of 56 universal single copy marker proteins (table S4) was used to place novel genomes into a phylogenetic tree together with a representative set of bacterial and archaeal reference genomes. Marker proteins were identified with hmmsearch (version 3.1b2, hmmer.org) using a specific hmm for each of the markers. For every protein, alignments were built with MAFFT (v7.294b, Katoh and Standley 2013) using the local pair option (mafft-linsi) and subsequently trimmed with trimAl 1.4^33^, removing sites for which more than 90 percent of taxa contained a gap. Query genomes lacking a substantial proportion of marker proteins (less than 28) or which had additional copies of more than one single-copy marker were removed from the data set. In total 17 of 29 Yellowstone hot spring genomes contained sufficient number of marker genes to be included in the tree. Single protein alignments were then concatenated resulting in an alignment of 51,239 sites. Maximum likelihood phylogenies were inferred with ExaML^34^ (version 3.0) using the GAMMA model for rate heterogeneity among sites and the LG substitution matrix^35^ and 300 non-parametric bootstraps. The resulting phylogenetic tree was visualized in ete3^36^.

### Comparison to shotgun metagenomics

Bulk genomic DNA were extracted from Yellowstone National Park hot spring samples using Qiagen’s blood and tissue kit using the protocol for DNA extraction from gram positive bacteria. Nextera V2 libraries were constructed and sequenced on Illumina’s Nextseq platform. 32.5 million and 51.4 million reads were obtained from Bijah Spring and Mound Spring samples and trimmed using the same parameters as the mini-metagenomic sequencing reads. Finally, assembly is performed using Megahit^37^ because SPAdes is not built to assemble shotgun metagenomic results. At the same time, we down sampled mini-metagenomic reads to the same depth as the shotgun metagenomic sequencing experiments and performed re-assembly using SPAdes.

### Analysis of genes involved in energy metabolism

From each genomic bin, ORFs assigned KO terms during the annotation process were mapped to all KEGG module involved in energy metabolism. For each KEGG module, we counted the number of KO terms extracted from a particular genome as a ratio of all KO terms present in the module. We did not normalize for genome size or completeness because doing so would artificially increase the importance of genes identified from smaller genomes.

### Assessment of genome completeness and abundance

Genome completeness and contamination (incorporation of contigs that may not belong to the genome) were quantified via CheckM^15^. Genome abundance was quantified using contig presence patterns. If more than 50% of the contigs from a particular bin were supported by at least one read in a sub-sample, we conclude that there was at least 1 cell in that sub-sample. For each genome, the number of sub-samples containing zero cells representing that particular genome was tabulated and Poisson distribution is used to calculate the most probable number of cells sampled in the microfluidic device. More specifically,

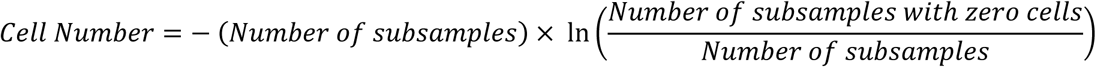

### Assessment of genome variation

SNPs (Single Nucleotide Polymorphism) were tabulated for all genomes from cells observed across all sub-samples by first aligning all reads to contigs in genome bins using samtools version 1.3 mpileup functionality with the –g flag^38^. Then, bcftools was used to call SNPs. We used several criteria to ensure confidence of observed SNPs. First, a SNP must have a quality score larger than 180 based on reads from all sub-samples. The results were not affected if we increase the threshold to higher numbers. Second, we required five reads to support each SNP location from a sub-sample in order to determine if the genome recovered from that sub-sample contains the dominant or alternate allele. Finally, we required that the alternate allele appear as the only allele in at least one sub-sample. If a sub-sample was determined to be heterozygous for a particular allele with high confidence, it was counted as one dominant and one alternate allele because it likely resulted from multiple cells from the same microfluidic chamber. Based on ORF predications, we then classified each SNP as noncoding, synonymous, or nonsynonymous.

## Data Availability

Fluidigm C1 IFC script is available at 
https://cn.fluidigm.com/c1openapp/scripthub Annotated contigs are available at 
https://img.jgi.doe.gov/mer/ under IMG Genome IDs 3300006068 and 3300006065.

Raw sequencing reads pending submission to NCBI SRA.

Software for contig assembly and binning can be made available upon request.

## Acknowledgements

The authors would like to acknowledge the Stanford Stem Cell Institute Sequencing Facility; Department of Energy (DOE) Joint Genome Institute’s (JGI) assembly and annotation teams; Rex Malmstrom (JGI); Danielle Goudeau (JGI); Anastasia Nedderton (Stanford); Jon Deaton (Stanford); and NPS staff at YNP for coordinating the sample collection process. This work is supported by DOE JGI Emerging Technologies Opportunities Program (ETOP) and Templeton Foundation. F.B.Y. is supported by SGF and NSF GRFP. P.C.B. is supported by the Burroughs Welcome Fund via a Career Award at the Scientific Interface. The work conducted by the U.S. Department of Energy Joint Genome Institute, a DOE Office of Science User Facility, is supported under Contract No. DE-AC02-05CH11231.

## Author Contributions

Conceptualization, F.B.Y., P.C.B., T.W., and S.R.Q.; Methodology, F.B.Y., F.S., T.W., M.A.H., and S.R.Q.; Software, F.B.Y., F.S.; Investigation, F.B.Y.; Writing, F.B.Y., F.S., T.W., and S.R.Q.; Supervision, M.A.H. and S.R.Q.

## Supplementary Files

**Fig. S1.**
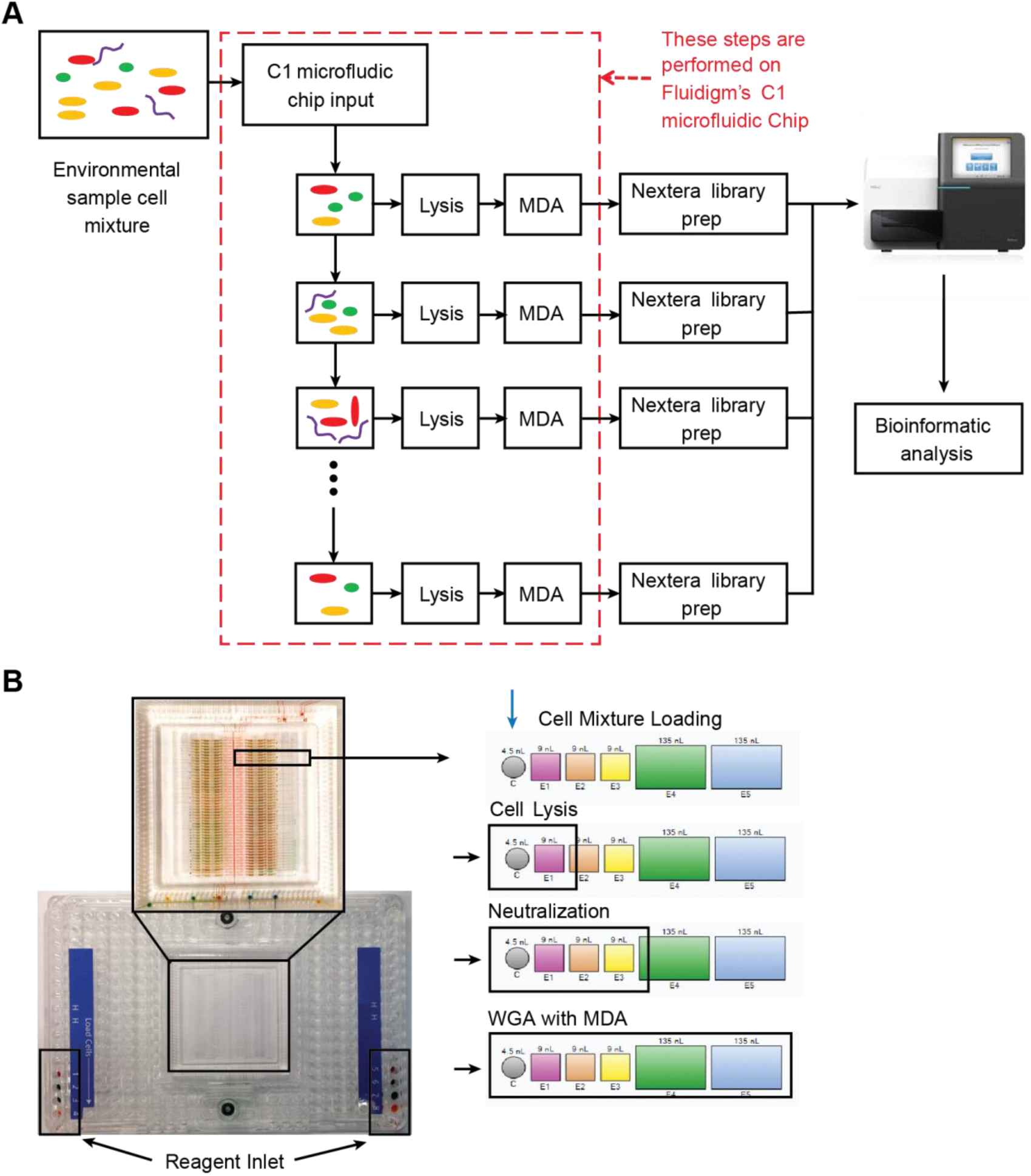
Schematic of the mini-metagenomic steps performed on the Fluidigm C1 microfluidic IFC. (A) Cells are distributed randomly into 96 chambers. Enzymatic lysis, DNA denaturation, and whole genome amplification via MDA occurs on chip. Amplified DNA is harvested for off-chip library preparations. (B) Structure of the Fluidigm IFC. The polydimethylsiloxane (PDMS) chip is located in the center of a plastic cartridge. Each chip contains 96 independent reaction systems. Each reaction system contains one capture chamber and five sequential reaction chambers of various sizes. Sequential reactions are carried out by incorporating additional chambers to provide volume for additional reagents. MDA is performed in all five chambers with a total volume of ~300 nL.

**Fig. S2.**
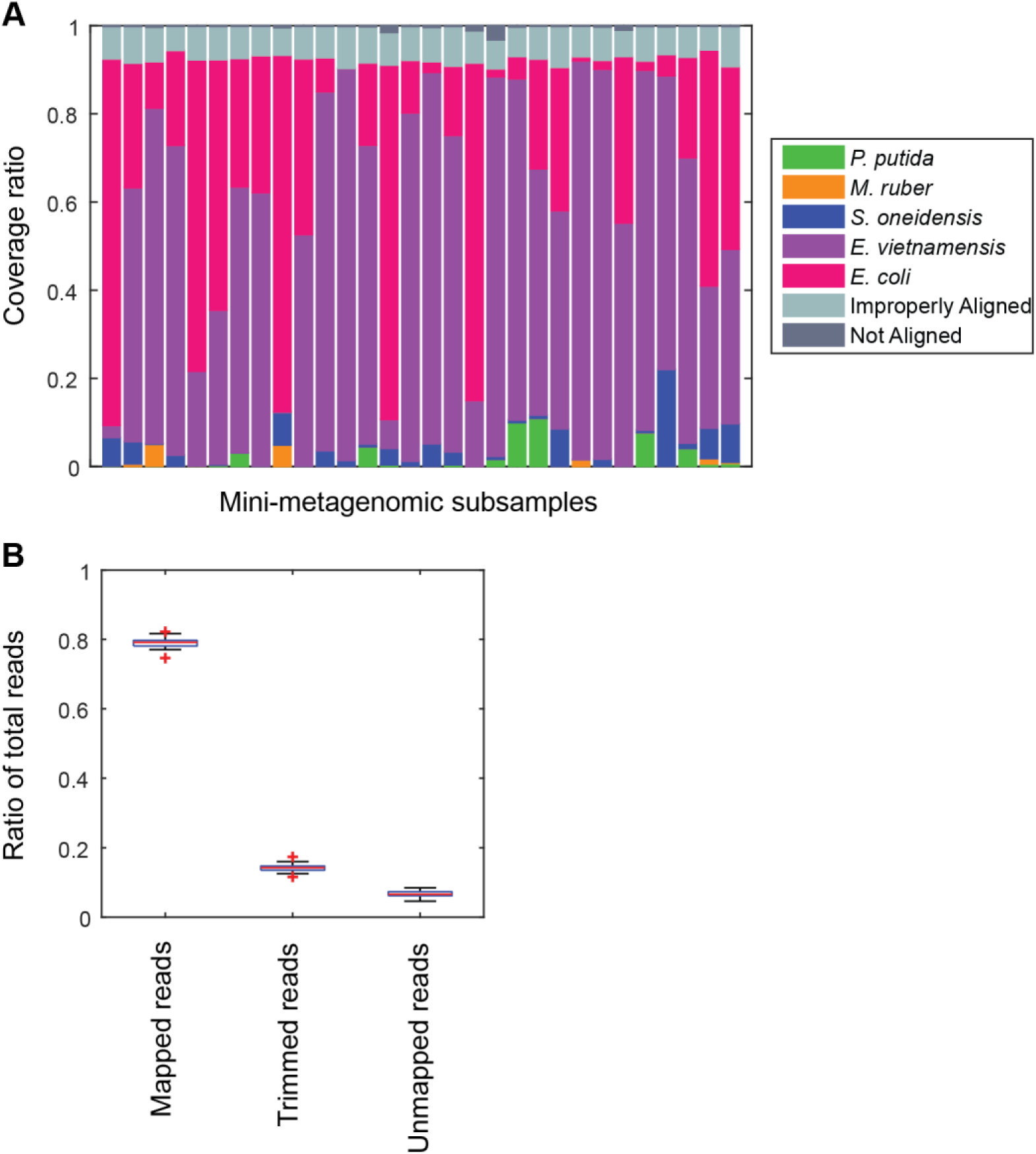
Performance of microfluidic-based mini-metagenomic amplification. A mock community is constructed and processed using the microfluidic-based mini-metagenomic pipeline. (A) Reads from each subsample are mapped back to known reference genomes. Over 90% of reads map uniquely to one of the 5 species. Improperly aligned reads are typically reads of poor sequencing quality or are too short to have a good alignment score. Less than 2% are unmapped. (B) Typically, 15% of raw sequencing reads from our mini-metagenomic pipeline are trimmed based on quality scores. 5% reads are unmapped, mostly represented by short library fragments. Less than 1% is chimeric reads.

**Fig. S3.**
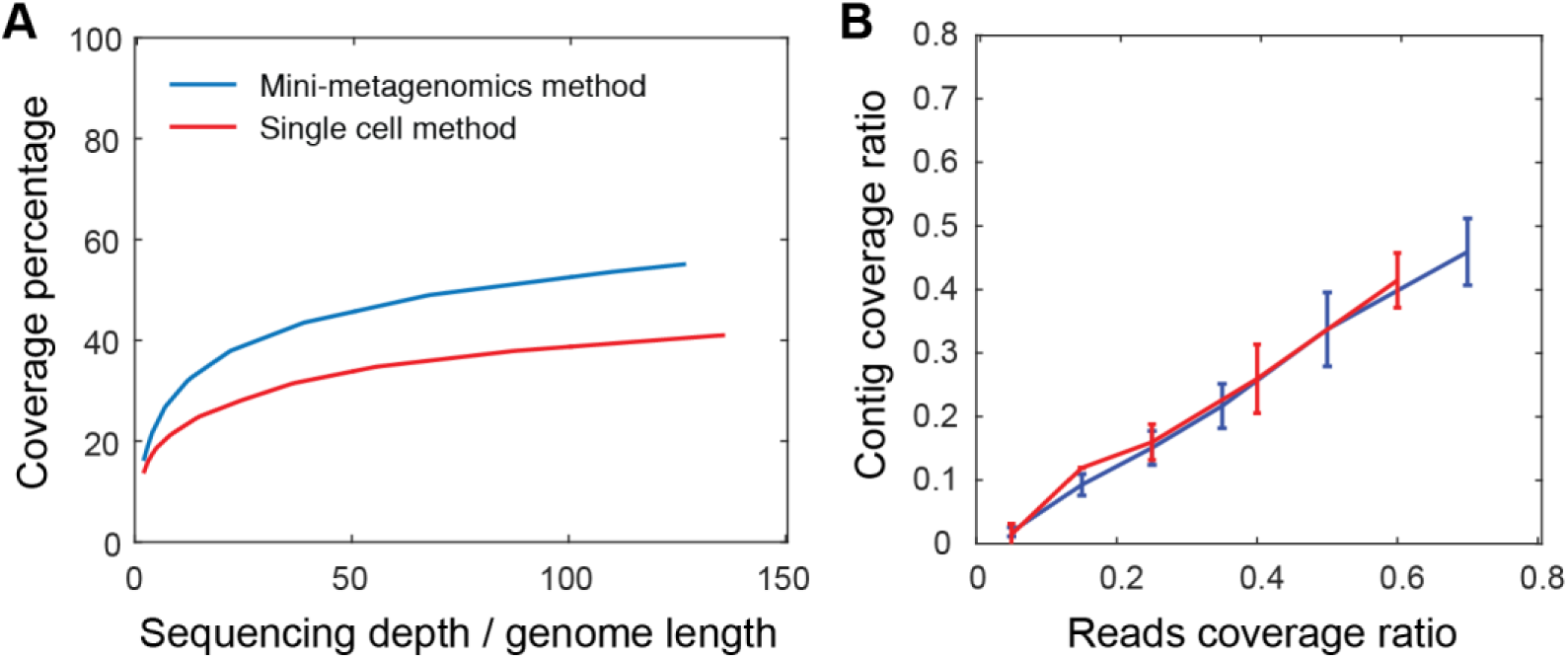
Mini-metagenomics performance on mock communities. (A) Compared to the single-cell method, mini-metagenomics showed a superior rarefaction curve. The percent genome coverage using the mini-metagenomic method is higher than using single-cell metagenomic methods for all sequencing depths. (B) Given a particular read coverage percentage, both mini-metagenomic and single-cell metagenomic methods yielded similar contig coverage ratio.

**Fig, S4.**
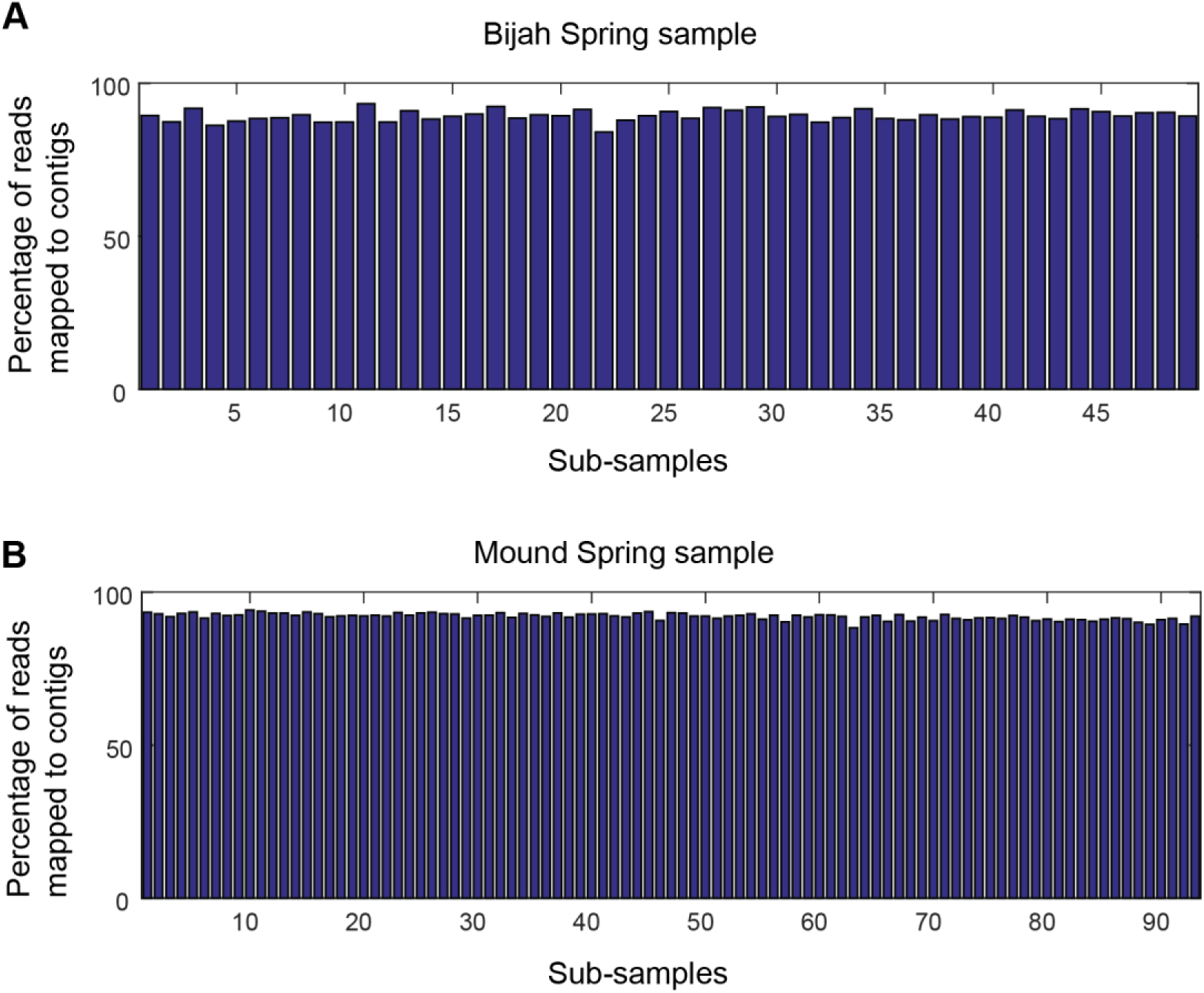
Approximately 90% of the reads are routinely incorporated into contigs 1 kbp or greater in every sub-sample for (A) Bijah and (B) Mound Spring samples.

**Fig. S5.**
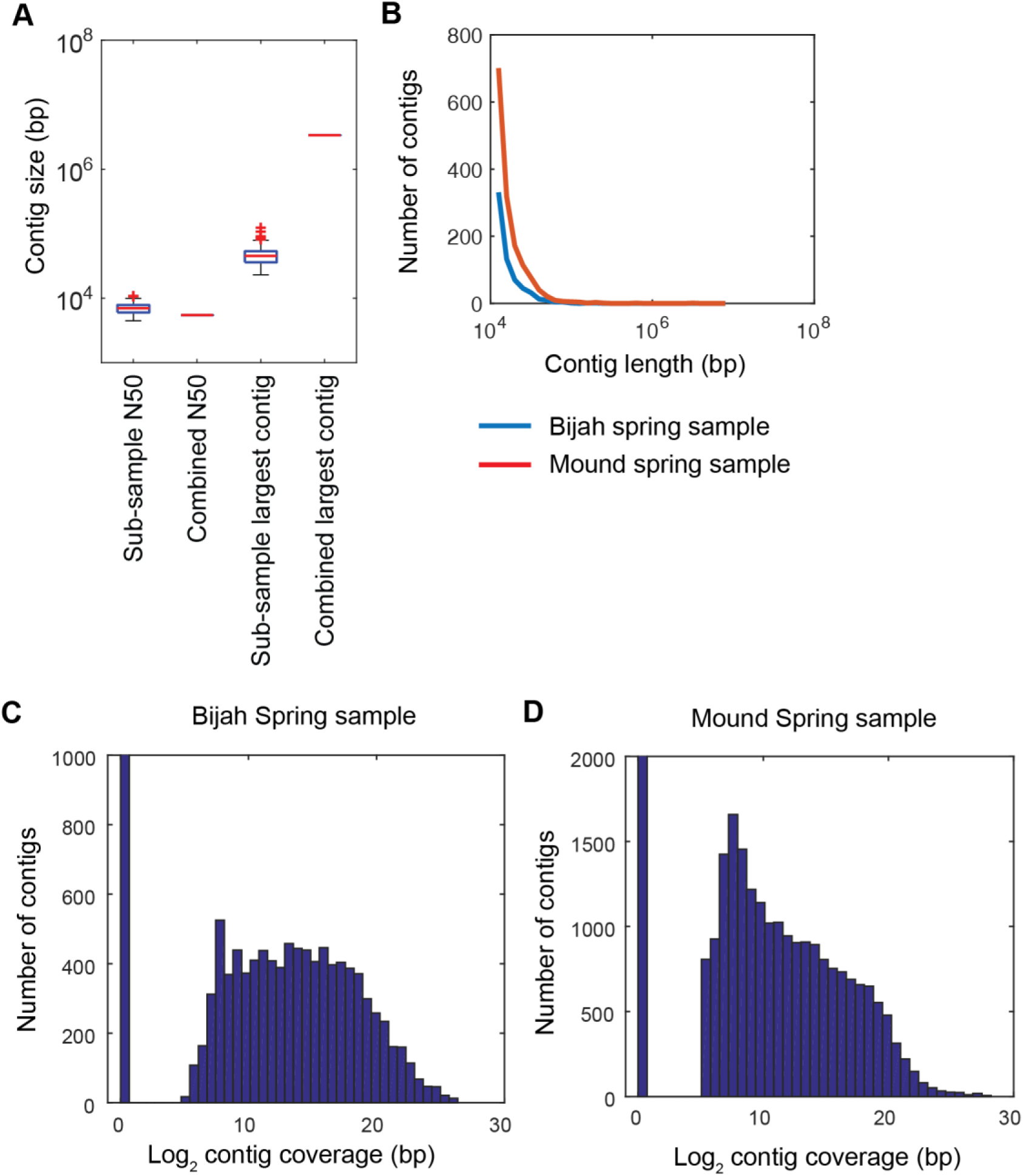
Contig statistics of Yellowstone National Park samples. (A) Sub-sample N50 and largest contig size in comparison with N50 and largest contig size for the combined assembly of all sub-samples reads. Assessment is done on contigs longer than 500 bps. Plot represents Mound Spring statistics. (B) Distribution of contig lengths for samples from Bijah (blue) and Mound (red) Springs. Plot shows only contigs greater than 10 kbp in length. Distribution of contig coverage from reads belonging to every sub-sample in (C) Bijah Spring and (D) Mound Spring samples. Coverage of 2^11^ bps is required for a contig to be designated as present in a sub-sample.

**Fig. S6.**
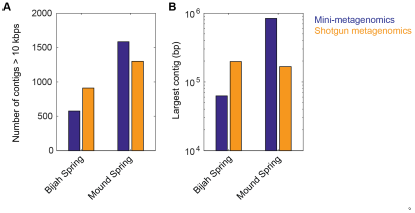
Comparison between contig statistics of mini-metagenomic and shotgun metagenomic assemblies. Mini-metagenomic reads from all sub-samples (49 for Bijah Spring and 93 for Mound Spring) were combined and randomly subsampled to the same depth as shotgun metagenomic sequencing depth of 32.5 million and 51.4 million reads, respectively. Mini-metagenomic reads were assembled using SPAdes ^31^ while Shotgun metagenomic reads were assembled using Megahit ^37^. Contigs over 10 kbps were tabulated using Quast ^39^. (A) Number of contigs over 10 kbps showing that shotgun metagenomics generated more contigs over 10 kbps in the Bijah Spring sample but less contigs in Mound Spring sample compared to mini-metagenomic assemblies. (B) Largest contig from shotgun metagenomic assembly was longer than the largest mini-metagenomic contig in the Bijah Spring sample but was shorter in the Mound Spring sample.

**Fig. S7.**
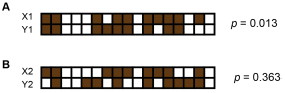
An example of computing *p* values using Fisher’s exact test using presence patterns of two contigs. Here, brown squares represent sub-samples in which the contig is present and white squares represent those from which the contig is missing. The computed *p* value using the equation

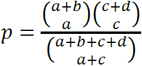 Where a = number of sub-samples where X is present and Y is present. b = number of sub-samples where X is absent and Y is present. c = number of sub-samples where X is present and Y is absent. d = number of sub-samples where X is absent and Y is absent. *p* represents the probability of incorrectly rejecting the null hypothesis that presence patterns of the two contigs X and Y occur randomly. (A) Presence patterns of X1 and Y1 are correlated because we obtain a small *p*. (B) We cannot reject the null hypothesis here due to the large *p* obtained.

**Fig. S8.**
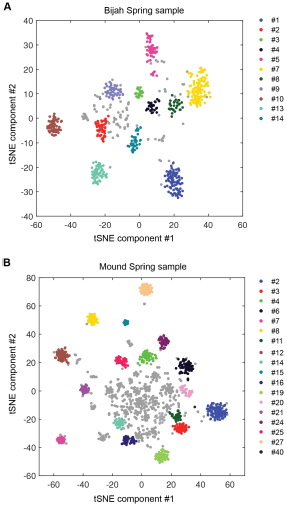
Cluster results applying DBscan to output of tSNE dimensional reduction based on pairwise *p* value matrix. Plots represent (A) Bijah Spring and (B) Mound Spring samples. Cluster numbers correspond to labels in Figure 2C, D. Gray contigs represent those that were not incorporated into a large enough contig cluster.

**Fig. S9.**
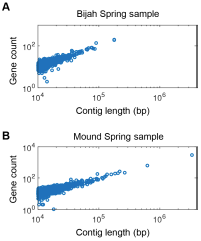
Gene count as a function of contig length for (A) Bijah and (B) Mound Spring samples. Only contigs larger than 10 kbp are shown. The linear relationship was consistent with little non-coding regions in bacterial genomes and increased the likelihood that these contigs represent biological entities instead of sequencing artifacts.

**Fig. S10.**
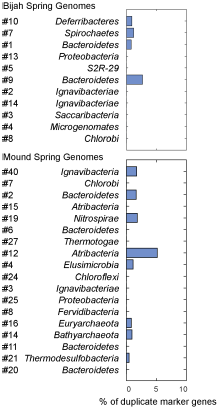
Degree of marker gene duplication in assembled genomes assessed using CheckM ^15^. All genomes except one have less than 2% marker gene duplication.

**Fig. S11.**
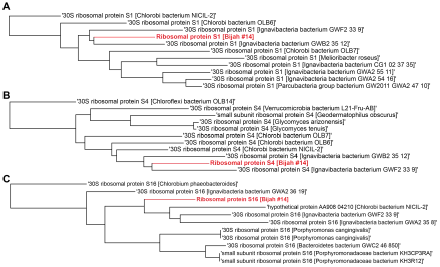
Single gene trees based on multiple sequence alignment of 10 most similar protein sequences for Bijah Spring genome #14 based on NCBI protein blast. Three longest ribosomal protein sequences identified in the genome were used including (A) ribosomal protein S1, (B) ribosomal protein S4, (C) ribosomal protein S16.

**Fig. S12.**
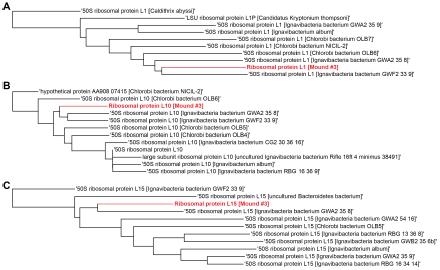
Single gene trees based on multiple sequence alignment of 10 most similar protein sequences for Mound Spring genome #3 based on NCBI protein blast. Three longest ribosomal protein sequences identified in the genome were used including (A) ribosomal protein L1, (B) ribosomal protein L10, (C) ribosomal protein L15.

**Fig. S13.**
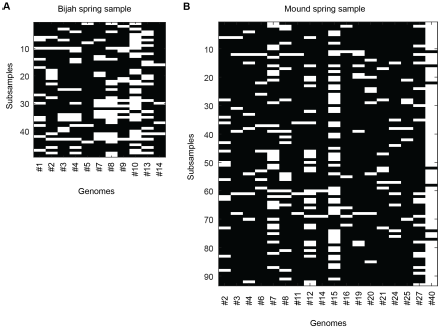
Abundance of each genome inferred from contig presence for (A) Bijah and (B) Mound Springs samples. White demonstrates the presence of at least one cell of a particular genome in a sub-sample. The total number of cells can be inferred using Poisson statistics.

**Fig. S14.**
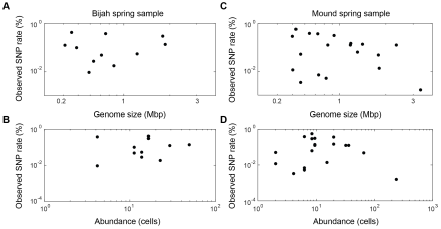
Observed SNP rate as a function of (A, C) genome size and (B, D) genome abundance for all genomes in (A, B) Bijah and (C, D) Mound Springs samples. SNP rate was quantified by summing all observed SNPs for a particular genome in all cells and normalizing to the total length of DNA sequenced from those cells. There did not seem to be any clear trend between SNP rate and genome size or abundance.

**Table S1:** Species used to construct mock bacterial communities

| Species name | GC content<br>(%) | Culture source | Assession numbers |
| --- | --- | --- | --- |
| Echinicola vietnamensis<br>KMM6221 | 44.7 | DSM 17526 | NC_019904.1 |
| Shewanella oneidensis MR1 | 45.9 | JGI | NC_004347.2 |
| Escherichia coli MG1655 | 50.8 | ATCC 700926 | NC_000913.3 |
| Pseudomonas putida F1 | 61.9 | ATCC 700007 | NC_009512.1 |
| Meiothermus ruber | 63.4 | DSM 1279 | NC_013946.1 |

**Table S2:** Information of hot spring samples from Yellowstone National Park

|  | Sample #1 | Sample #2 |
| --- | --- | --- |
| NPS study number | YELL_05788 | YELL_05788 |
| Sample area | Mammoth Norris Corridor | Lower Geyser Basin |
| Location name | Bijah Spring | Mound Spring |
| Sample type | Sediment | Sediment |
| Collection time | September 11, 2009 5:00pm | September 14, 2009 2:25pm |
| Location | 44.761133 N, 110.730900 W | 44.564833 N, 110.859933 W |
| Location type | Hot spring | Hot spring |
| Temperature (°C) | 76 | 90 |
| pH | 7.0 | 9.0 |

**Table S3:** Yellowstone sample sequencing and assembly statistics

|  | Sample #1 | Sample #2 |
| --- | --- | --- |
| IMG genome ID | 3300006068 | 3300006065 |
| Number of molecules sequenced | 120.5 M | 133.1 M |
| Number of contigs (>10kbp) | 643 | 1474 |
| Number of subsamples analyzed | 49 | 93 |

**Table S4:** COG terms used for marker gene based phylogenetic tree

| COG id | Gene name |
| --- | --- |
| COG0013 | alanyl-tRNA_synthetase |
| COG0016 | phenylalanyl-tRNA_synthetase_alpha_subunit |
| COG0018 | arginyl-tRNA_synthetase |
| COG0048 | ribosomal_protein_S12 |
| COG0049 | ribosomal_protein_S7 |
| COG0051 | ribosomal_protein_S10 |
| COG0052 | ribosomal_protein_S2 |
| COG0060 | isoleucyl-tRNA_synthetase |
| COG0072 | phenylalanyl-tRNA_synthetase_beta_subunit |
| COG0080 | ribosomal_protein_L11 |
| COG0081 | ribosomal_protein_L1 |
| COG0085 | DNA-directed_RNA_polymerase_beta_subunit (RpoB) |
| COG0086 | DNA-directed_RNA_polymerase_beta_prime_subunit (RpoC) |
| COG0087 | ribosomal_protein_L3 |
| COG0088 | ribosomal_protein_L4 |
| COG0089 | ribosomal_protein_L23 |
| COG0090 | ribosomal_protein_L2 |
| COG0091 | ribosomal_protein_L22 |
| COG0092 | ribosomal_protein_S3 |
| COG0093 | ribosomal_protein_L14 |
| COG0094 | ribosomal_protein_L5 |
| COG0096 | ribosomal_protein_S8 |
| COG0097 | ribosomal_protein_L6P |
| COG0098 | ribosomal_protein_S5 |
| COG0099 | ribosomal_protein_S13 |
| COG0100 | ribosomal_protein_S11 |
| COG0102 | ribosomal_protein_L13 |
| COG0103 | ribosomal_protein_S9 |
| COG0127 | Xanthosine_triphosphate_pyrophosphatase |
| COG0130 | Pseudouridine_synthase |
| COG0164 | ribonuclease_HII |
| COG0172 | seryl-tRNA_synthetase |
| COG0184 | ribosomal_protein_S15P |
| COG0185 | ribosomal_protein_S19 |
| COG0186 | ribosomal_protein_S17 |
| COG0193 | peptidyl-tRNA_hydrolase |
| COG0197 | ribosomal_protein_L16 |
| COG0198 | ribosomal_protein_L24 |
| COG0200 | ribosomal_protein_L15 |
| COG0201 | preprotein_translocase_subunit_SecY |
| COG0202 | DNA-directed_RNA_polymerase_alpha_subunit (RpoA) |
| COG0216 | protein_chain_release_factor_A |
| COG0233 | ribosome_recycling_factor |
| COG0244 | ribosomal_protein_L10 |
| COG0255 | ribosomal_protein_L29 |
| COG0256 | ribosomal_protein_L18 |
| COG0343 | queuine/archaeosine_tRNA-ribosyltransferase |
| COG0481 | membrane_GTPase_LepA |
| COG0495 | leucyl-tRNA_synthetase |
| COG0504 | CTP_synthase |
| COG0519 | GMP_synthase_PP-ATPase_domain |
| COG0532 | translation_initiation_factor_2 |
| COG0533 | metal-dependent_proteases_with_possible_chaperone_activity |
| COG0541 | signal_recognition_particle_GTPase |
| COG0691 | tmRNA-binding_protein |
| COG0858 | ribosome-binding_factor_A |

